# AAK1 inhibits WNT signaling by promoting clathrin-mediated endocytosis of LRP6

**DOI:** 10.1101/258632

**Authors:** Megan J. Agajanian, Matthew P. Walker, Alison D. Axtman, Roberta R. Ruela-de-Sousa, Alex D. Rabinowitz, David M. Graham, Meagan Ryan, D. Stephen Serafin, James M. Bennett, Rafael M. Couñago, David H. Drewry, Jonathan M. Elkins, Carina Gileadi, Opher Gileadi, Paulo H. Godoi, Nirav Kapadia, Susanne Müller, André S. Santiago, Fiona J. Sorrell, Carrow I. Wells, Oleg Fedorov, Timothy M. Willson, William J. Zuercher, Michael B. Major

**Affiliations:** Lineberger Comprehensive Cancer Center, University of North Carolina at Chapel Hill, Chapel Hill, North Carolina, USA; Department of Pharmacology, University of North Carolina at Chapel Hill, Chapel Hill, North Carolina, USA; Structural Genomics Consortium, UNC Eshelman School of Pharmacy, University of North Carolina at Chapel Hill, Chapel Hill, NC, USA; Division of Chemical Biology and Medicinal Chemistry, UNC Eshelman School of Pharmacy, University of North Carolina at Chapel Hill, Chapel Hill, NC, USA; Structural Genomics Consortium, Universidade Estadual de Campinas — UNICAMP, Campinas, SP, Brazil; Department of Cell Biology and Physiology, University of North Carolina at Chapel Hill, Chapel Hill, North Carolina, USA; Structural Genomics Consortium and Target Discovery Institute, Nuffield Department of Clinical Medicine, University of Oxford, Old Road Campus Research Building, Oxford, OX3 7DQ, UK.; Structural Genomics Consortium, Buchmann Institute for Molecular Life Sciences, Goethe University Frankfurt, Frankfurt am Main, Germany; Department of Computer Science, University of North Carolina at Chapel Hill, Chapel Hill,North Carolina 27599, USA

**Keywords:** WNT signaling, endocytosis, gain-of-function screen, AAK1, clathrin, kinase

## Abstract

β-catenin-dependent WNT signal transduction governs normal development and adult tissue homeostasis. Inappropriate pathway activity mediates a vast array of human diseases, including bone density disorders, neurodegeneration and cancer. Although several WNT-directed therapeutics are in clinical trials, new targets, compounds and strategies are needed. We performed a gain-of-function screen of the human kinome to identify new druggable regulators of β-catenin-dependent transcription. We found that over-expression of the AP2 Associated Kinase 1 (AAK1) strongly inhibited WNT signaling. Reciprocally, silencing of AAK1 expression or pharmacological inhibition of AAK1 kinase activity using a new, selective and potent small molecule inhibitor activated WNT signaling. This small molecule is a cell active dual AAK1/BMP2K inhibitor that represents the best available tool to study AAK1-dependent signaling pathways. We report that AAK1 and the WNT co-receptor LRP6 physically co-complex and that AAK1 promotes clathrin-mediated endocytosis of LRP6. Collectively, our data support a WNT-induced negative feedback loop mediated by AAK1-driven, clathrin-mediated endocytosis of LRP6.

**Summary Statement:** A gain-of-function screen of the human kinome revealed AAK1 as a negative regulator of WNT signaling. We show that AAK1 promotes clathrin-mediated endocytosis of LRP6, resulting in downregulation of WNT signaling. We use a new selective and potent AAK1/BMP2K small molecule probe to validate our findings.

## Introduction

β-catenin-dependent WNT signal transduction is an evolutionarily conserved pathway that instructs cell and tissue-specific differentiation and proliferation programs. It is both required for normal development and maintenance of adult tissue homeostasis^1^. Like many developmental signaling pathways, aberrant WNT signaling contributes fundamentally to human disease, including cancer, bone density disorders and neurodegeneration^1^. The regulation of the WNT/β-catenin signaling axis is often described in three stages: proximal signaling, β-catenin destruction and nuclear effectors of transcription^2^. In the absence of WNT ligand, the LRP5/6 (Low-density lipoprotein receptor related protein 5/6) and FZD (Frizzled) family of WNT receptors remain dissociated. Within the cytosolic compartment, a multi-protein β-catenin destruction complex comprises AXIN1/2, APC (Adenomatous polyposis coli), CK1α(Casein kinase 1α), and GSK3β (Glycogen synthase kinase 3β). CK1α and GSK3β sequentially phosphorylate β-catenin, resulting in its ubiquitylation by β-TRCP and ultimately proteasomal degradation^3–6^. Extracellular WNT ligand physically couples LRP5/6 and FZD receptors, leading to phosphorylation-dependent recruitment of the AXIN1/2, APC, GSK3β and DVL (Dishevelled1/2/3) proteins^7^. This complex, referred to as the LRP6 signalosome, transiently suppresses β-catenin phosphorylation by GSK3β^7^. Hence, WNT ligand transiently inactivates the destruction complex to promote cytoplasmic accumulation of β-catenin, resulting in its translocation to the nucleus, where it binds to members of the TCF/LEF family of transcription factors to modulate tissue-specific transcriptional programs and cell phenotypes^8^.

Through a gain-of-function screen of the kinome, we discovered that the AAK1 (Adaptor Associated Kinase 1) kinase negatively regulates β-catenin-dependent WNT signaling by promoting clathrin-mediated endocytosis (CME) of LRP6. CME is a complex and multi-step process that removes lipids and transmembrane proteins from the plasma membrane. The assembly polypeptide 2 (adaptor-related protein 2; AP2) protein complex first recognizes and binds sorting signals on the intracellular domains of transmembrane cargo proteins^9,10^. Activation of the AP2 machine occurs allosterically through binding cargo, clathrin and PIP2 (Phosphatidylinositol 4,5-bisphosphate)^11^. This activation coincides with a conformational shift in AP2 from “closed” to “open”. Open/active AP2 is required for continued clathrin polymerization, cargo recruitment, and ultimately the maturation of a nascent clathrin-coated pit to a stable, successful endocytic event^10–14^. In addition to PIP2, cargo and clathrin, activation of the AP2 complex is promoted by phosphorylation of the AP2M1 subunit by AAK1^10–14^. Importantly, AAK1-dependent phosphorylation of AP2M1 is promoted by clathrin assembly^13^, which constitutes a positive feed-forward loop to promote pit maturation. Once mature, cargo-loaded clathrin-coated pits evolve to endosomal vesicles through dynamin-mediated scission from the plasma membrane^15^.

Numerous studies support a role for CME in both positive and negative regulation of WNT signaling, though in either scenario a specific role for AAK1 has not been reported^16–20^. First, clathrin, AP2 and PIP2 are required for WNT-induced LRP6 phosphorylation, signalosome formation and subsequent WNT signaling^7,19^. AP2 and clathrin recruitment to the LRP6 signalosome is induced by WNT ligand, presumably via WNT-induced PIP2 production^19^. Indeed, WNT ligand induces a gradual accumulation of PIP2 over a 4-hour after treatment with WNT ligand^21^. Together, these studies suggest that clathrin-associated machinery is required for WNT-induced signalosome formation. Second, others have reported that CME of LRP6 downregulates WNT signaling^20^. For example, disabled homolog 2 (DAB2) drives LRP6 toward clathrin-mediated endocytosis and represses WNT signaling^18^. Additionally, phosphorylation of LRP6 on cytoplasmic tyrosine residues drives CME of LRP6, resulting in decreased WNT signaling^22^.Third, De Robertis and colleagues described a mechanism by which the endocytosed signalosome is incorporated into multivesicular bodies (MVBs). Here, sequestration of GSK3β in MVBs suppresses β-catenin phosphorylation, thus potentiating signaling^23^. Last, Gammons and colleagues describe how signalosome incorporation into clathrin-coated pits allows for DVL polymerization via domain swapping, an event which promotes WNT signaling activation^24^. All together, these studies establish CME as a major regulatory feature in WNT signaling, although contextual and temporal considerations remain largely unresolved.

Cross-referencing our gain-of-function screen of the kinome with previously published loss-of-function screens, we identify AAK1 as a negative regulator of WNT-induced β-catenin transcription. Gain- and loss-of-function studies reveal that AAK1 negatively regulates WNT signaling by promoting CME of LRP6. We also report and employ a novel potent and selective cell active dual AAK1/BMP2K inhibitor, which potentiates WNT signaling. Full characterization of this small molecule supports that it is the best available tool to interrogate AAK1 biology. Unexpectedly, we discovered that WNT induces phosphorylation of AP2M1 by AAK1 in a temporally delayed fashion at 8-10 hours post-WNT3A treatment. Therefore, we propose a modified model whereby WNT-driven CME promotes a negative feedback mechanism to limit pathway longevity. More broadly, this work contributes to the ongoing discussion of endocytosis within WNT signaling by solidifying a role for the AAK1 kinase in removing LRP6 from the cell plasma membrane.

## Results

### eIdentification of AAK1 as a novel repressor of WNT signaling

Thirty years of research on the WNT pathway has revealed its core components and basic mechanics. That said, recent improvements in genetic screening technologies and protein mass spectrometry continue to identify novel modifiers of WNT pathway activity. We, along with others, have reported genome-wide gain- and loss-of-function genetic screens of the WNT pathway, illuminating new regulators (eg. FOXP1, USP6, BRAF^V600E^) and their mechanisms of WNT control^25–27^. Here, we focused on the human kinome to determine which kinases when over-expressed activate or inhibit WNT3A-driven β-catenin transcription. We cloned 387 kinase open reading frames (ORF) into a lentiviral expression vector before transient over-expression in HEK293T cells carrying the β-catenin-activated Firefly luciferase reporter (BAR) and a constitutively expressed *Renilla (Ren)* luciferase. Eighteen hours after kinase transfection, cells were treated with WNT3A conditioned media (CM) for an additional 16 hours before lysis and BAR/*Ren* reporter quantitation (**Supplementary Table 1**). ANKRD6/Diversin and CTNNB1/β-catenin served as negative and positive controls, respectively. Five kinases (AAK1, ADCK1, ADCK2, MAST1 and TGFBR3) were validated by low throughput reporter assays in HEK293TBAR/*Ren* cells (**Fig. 1a**). Comparative analysis of this kinome gain-of-function screen in HEK293T cells with two previously published siRNA-based loss-of-function screens in HT1080 sarcoma cells and A375 melanoma cells revealed a single common protein: AP2 Associated Kinase 1 (AAK1) (**Supplementary Table 1**)^25,27^. Because of the well-established functional connections between AAK1 and CME, and the emerging data on CME in governing WNT pathway dynamics, we sought to understand how AAK1 negatively regulates the WNT pathway.

**Figure 1.**
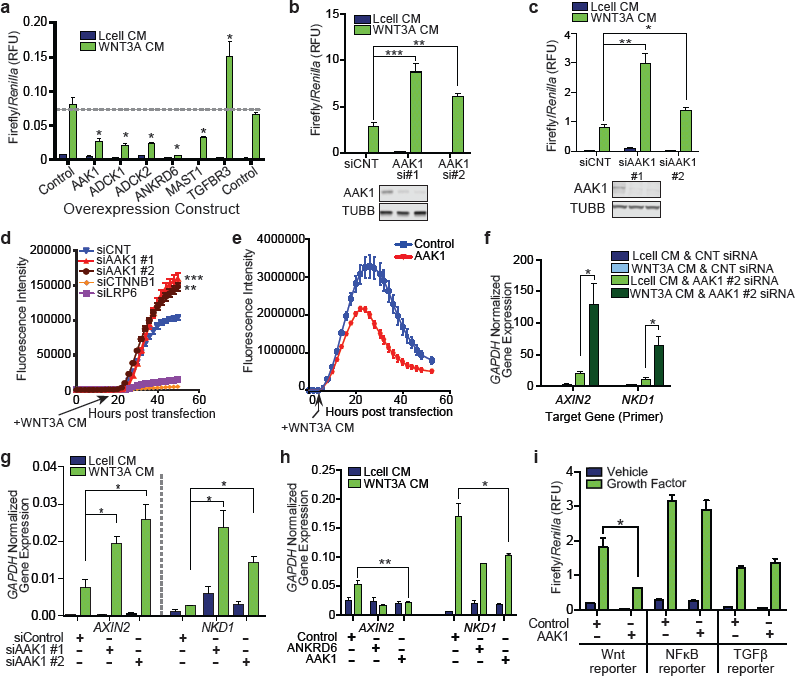
Gain-of-function kinome screen reveals AAK1 as a repressor of WNT signaling. **(a)** HEK293T-B/R cells were transfected with the indicated construct for 24 hours. Cells were then treated for 16 hours with WNT3A or Lcell CM. Bars represent average Firefly/*Ren* relative fluorescence units (RFU) from 3 technical replicates. **(b,c)** Luciferase assay of HT1080 **(c)** or **(c)** stable B/R cells transfected with either control or AAK1 siRNA constructs for 56 hours. Cells were then treated with either Lcell or WNT3A CM for 16 hours. Bars represent average Firefly/*Ren* RFU from 3 technical replicates. Western blot analysis illustrates knock down efficiency of two independent AAK1 siRNAs. **(d)** IncuCuyte imaging of HT1080 cells stably expressing a BAR-mCherry fluorescent reporter transiently transfected with indicated siRNA construct. WNT3A CM was added at 18 hours, then cells were imaged for 50 hours post-transfection. Graph represents data points averaged across 4 technical replicates. Live cell imaging of HT1080 cells stably expressing a BAR-mCherry fluorescent reporter transiently transfected with the indicated expression construct, AAK1 or Flag Control. WNT3A CM was added at 8 hours, and cells were monitored for an additional 56 hours. Data represent the average of 4 technical replicates. **(f,g)** qPCR analysis of *AXIN2* and *NKD1* in HEK293T **(f)** or HT1080 **(g)** cells 72 hours after transfection with the indicated siRNA. Cells were treated with WNT3A CM for 6 hours prior to harvest. Bars represent average glyceraldehyde-3-phosphate dehydrogenase (*GAPDH*)-normalized gene expression across 3 technical replicates. **(h)** qPCR analysis of *AXIN2* (left) and *NKD1* (right) in HEK293T cells transfected with over-expression construct for 24 hours, then treated with WNT3A CM for 6 hours prior to harvest. Bars represent average *GAPDH*-normalized gene expression across 3 technical replicates. **(i)** Luciferase assay of HEK293T cells transfected with indicated pathway-specific Firefly luciferase reporter constructs and expression constructs prior to a 16 hour treatment with recombinant human (rh) WNT3A (200ng/mL), rhTNFα (200ng/mL) or rhTGFβ1 (200ng/mL). Bars represent average Firefly/*Ren* RFU from 3 technical replicates. All panels:*** p<0.0005, ** p<0.005, * p<0.05. Data are representative of biological triplicates, unless otherwise noted. For complete statistics, see methods and materials.

To validate and extend the discovery of AAK1 as a WNT inhibitor, we tested: 1) whether siRNA-mediated silencing of AAK1 activated β-catenin-driven transcription, 2) the cell-type specificity of the AAK1-WNT phenotype, 3) whether AAK1 regulated the expression of endogenous β-catenin target genes, and 4) whether AAK1 affected the activity of non-WNT signaling pathways. First, in agreement with AAK1 over-expression blocking WNT signaling (**Fig. 1a**), siRNA silencing of AAK1 using two non-overlapping siRNAs increased BAR expression in HT1080 fibrosarcoma cells and RKO colon cancer cells (**Fig. 1b,c**). To visualize reporter expression in real time, we silenced AAK1 in HT1080 cells carrying a BAR-mCherry reporter. Quantitation of mCherry fluorescence confirmed that AAK1 knockdown activated the BAR reporter (**Fig. 1d**), while AAK1 over-expression suppressed BAR activity (**Fig. 1e**). Third, to rule out potential reporter-based artifacts, we quantified the expression of two WNT target genes after AAK1 perturbation. AAK1 knockdown increased RNA expression of *AXIN2* and *NKD1* in both HEK293T and HT1080 cells (**Fig. 1f,g**). Conversely, AAK1 over-expression led to decreased RNA expression of *AXIN2* and *NKD1* in HEK293T cells (**Fig. 1h**). Fourth, because of its established roles in CME, AAK1 might broadly regulate other signaling cascades. AAK1 over-expression did not affect TNFα-driven NFκB reporter activity or TGFβ-driven SMAD reporter activity (**Fig. 1i**). Together, these data establish that AAK1 negatively regulates WNT signaling in cells derived from multiple tissue types. Importantly and consistent with its established role in CME, AAK1 did not impact β-catenin transcriptional activity in the absence of exogenous WNT3A stimulation.

### Discovery of a potent and selective inhibitor of AAK1

Indazole compound SB-742864 was previously identified as a semi-selective inhibitor of AAK1 (*in vitro* IC_50_ = 220 nM)^28^. Optimization of SB-742864 via iterative medicinal chemistry optimization led to compound 25A (**Fig. 2a**), a potent and selective inhibitor of AAK1 and the closely related kinase, BMP2K. Further optimization to improve the selectivity led to SGC-AAK-1 (**Fig. 2a**), a chemical probe for AAK1 and BMP2K. A closely related molecule, SGC-AAK1-1N (referred to as 34A), which is devoid of activity on AAK1 or BMP2K, was identified for use as a negative control compound. Full details of the chemistry program and analysis of the structure-activity relationships will be published elsewhere. Compounds 25A and SGC-AAK-1, but not 34A, both selectively inhibited AAK1 and BMP2K over the other members of the subfamily, Numb-associated kinases GAK and STK16, in a TR-FRET binding displacement assay (**Fig. 2b, Supplementary Fig. 1a**), with *K*_i_ values against AAK1 of 8 nM and 9 nM respectively for 25A and SGC-AAK1-1 and 2 µM for negative control 34A. Both 25A and SGC-AAK1-1 showed good selectivity over a panel of 406 wild-type protein kinases at 1 µM concentration (**Fig. 2c, Supplementary Fig. 1b, Supplementary Table 2**) with only a small number of other kinases inhibited with *K*_D_ values within 30x of those of AAK1. Isothermal titration calorimetry measured *K*_D_ values for SGC-AAK1-1 of 120 nM and 490 nM against AAK1 and BMP2K respectively and showed that binding had favorable enthalpy and unfavorable entropy (**Fig. 2d, Supplementary Fig. 1c, Supplementary Table 3**). Attempts to co-crystallise either inhibitor with AAK1 were unsuccessful, but it was possible to obtain a co-crystal structure of 25A with BMP2K to 2.41 Å resolution (**Fig. 2e, Supplementary Fig. 1d,e,f, Supplementary Table 4**). The structure revealed 25A bound in the ATP site, making three hydrogen bonds to the kinase hinge region and two further hydrogen bonds via its sulfonamide oxygens to Gln137 and Asn185, while the central hydrophobic portion of the inhibitor packs against the gatekeeper residue Met130. To test that the inhibitors engage AAK1 and BMP2K in live cells we used NanoBRET assays (Promega)^29^, in which human cells were transfected with a plasmid expressing the full-length kinase gene fused to Nanoluc luciferase. In the presence of a fluorescent-labelled ATP-competitive tracer compound a BRET signal was measured. Displacement of the tracer by the inhibitor being tested demonstrated target engagement, and linear regression of the IC_50_ values at multiple tracer concentrations yielded the IC_50_ in the absence of tracer. 25A and SGC-AAK-1 bound to AAK1 in HEK293T cells with IC_50_ values of around 240 nM, and more weakly to BMP2K, with IC_50_ values of 600 nM and 1.5 µM respectively (**Fig. 2f,g,h, Supplementary Fig. 1g**). To demonstrate that binding to AAK1 inhibited its enzymatic activity in cells, phospho-AP2M1 (Thr156) levels were observed by Western blotting after 2 hour treatment of HEK293T cells with serially diluted SGC-AAK1-1, resulting in a dose-dependent decrease of pAP2M1 starting below 1 µM in agreement with the NanoBRET data (**Fig. 2i**).

**Figure 2.**
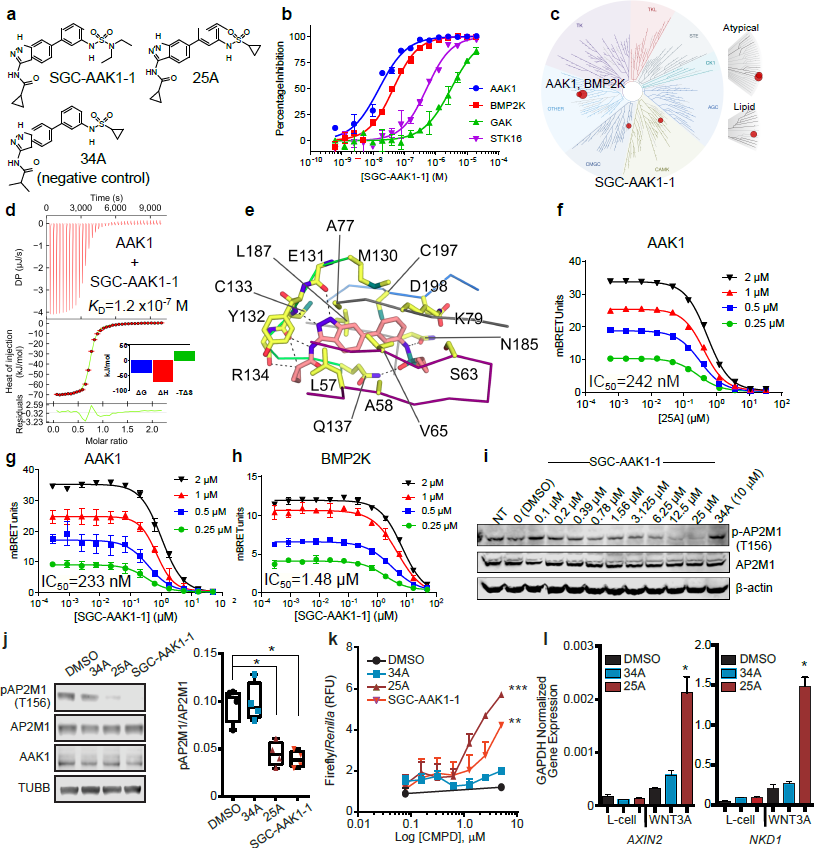
SGC-AAK1-1 is a potent and specific inhibitor of AAK1 and BMP2K that is active in cells. **(a)** Chemical structures of the AAK1/BMP2K chemical probe SGC-AAK-1, the related AAK1/BMP2K inhibitor 25A, and the negative control compound 34A that has a similar chemical structure but does not inhibit AAK1 or BMP2K. **(b)** SGC-AAK-1 selectively bound AAK1 and BMP2K with a more than 30-fold difference in Ki in a TR-FRET binding displacement assay over the related kinases GAK and STK16. 16-point dose-response curves were measured in duplicate. **(c)** SGC-AAK1-1 has good selectivity over the human kinome. SGC-AAK1-1 was used at 1 µM concentration. Kinases for which SGC-AAK1-1 has a KD < 100 nM are marked with a large red circle and 100 nM < KD < 1.0 µM with a smaller red circle. **(d)** Isothermal titration calorimetry confirmed a dissociation constant (KD) of 120 nM for binding of SGC-AAK1-1 **(e)** A co-crystal structure of BMP2K (2.41 Å) bound to 25A revealed the binding mode of the inhibitor in the ATP site. **(f,g)** NanoBRET cellular target engagement assays in HEK293T cells showed that 25A **(f)** and SGC-AAK1-1 **(g)** both entered cells and directly bound to AAK1 and displaced a fluorescent tracer molecule from the ATP site. Cells were treated with serially diluted inhibitors in the presence of four different concentrations of a NanoBRET tracer molecule, and IC_50_ values were calculated from a linear interpolation of values at each tracer concentration to obtain the predicted IC_50_ in the absence of tracer. Measurements were made with three technical replicates. **(h)** Binding of SGC-AAK1-1 to BMP2K in cells was weaker than to AAK1 with IC_50_ > 1 µM. **(i)** SGC-AAK1-1 inhibited the phosphorylation of the AP2M1 subunit at Thr156 in HEK293T cells in a concentration-dependent manner. HEK293T cells were treated with inhibitors for 2 hours before Western blot analysis. **(j)** Representative Western blot of HT1080 cells treated with indicated compounds (2.5 µM) for one hour (left). Box and whisker plot represents quantification of four biological replicate experiments using LiCOR software (right). Data is representative of 4 biological replicates. **(k)** BAR luciferase assay of HT1080-B/R stable cells treated with WNT3A CM and the indicated dose of compound for 16 hours. Data are averaged over 3 technical replicates. **(l)** qPCR analysis of *AXIN2*from HT1080 cells treated with indicated compound (1.25 µM) for 12 hours in the presence of Lcell or WNT CM and normalized. Graph represents analysis averaged across 3 technical replicates. All panels:*** p<0.0005, ** p<0.005, * p<0.05. Data are representative of biological triplicates, unless otherwise noted. For complete statistics, see methods and materials.

To confirm and extend these data, we next tested SGC-AAK1-1 and 25A in a series of experiments to evaluate its potency of AAK1 inhibition in cells and regulation of WNT signaling. First, we treated HT1080 cells with SGC-AAK1-1 and evaluated AAK1-dependent phosphorylation of AP2M1. SGC-AAK1-1, along with 25A, both significantly reduced phosphorylation of AP2M1 (T156), compared to controls, DMSO and 34A (**Fig. 2j**). Second, treatment with SGC-AAK1-1 or 25A activated WNT-driven BAR activity compared to controls (DMSO and 34A) in a dose-dependent manner (**Fig. 2k**). Importantly, 25A also significantly upregulated WNT target genes *AXIN2* and *NKD1* in HT1080 cells, as compared to DMSO control and 34A (**Fig. 2l**). Together, these results indicate that 25A and SGC-AAK1-1 block AAK1 kinase activity, resulting in increased WNT/β-catenin signaling.

### AAK1 promotes CME of LRP6 to inhibit WNT signaling

Several recent studies report that CME negatively regulates WNT signaling by reducing the presence of the receptor complex on the plasma membrane^18,20,22^. Given the well-established role of AAK1 in phosphorylating AP2M1 to facilitate CME^10,12–14^, our discovery of AAK1 as a negative regulator of WNT signaling is not entirely unexpected. That said, AAK1 has not previously been shown to regulate WNT signaling, and certainly, the molecular mechanisms governing CME within the WNT pathway and its homeostasis are not well-defined, with several remaining questions and conflicting studies^16–18,20,22–24^. To confirm that AAK1 functions within membrane proximal steps of WNT signaling, we stabilized β-catenin protein levels with a GSK3β inhibitor (CHIR99021) following either AAK1 over-expression or siRNA silencing. CHIR99021-mediated activation of the BAR reporter was not affected by AAK1 depletion or over-expression (**Fig. 3a**). As expected, β-catenin knockdown blocked BAR activation by AAK1 silencing (**Supplementary Fig. 2a**). These results indicate that AAK1 functions upstream of β-catenin and the destruction complex. Next, the role of endocytosis in AAK1-mediated repression of WNT signaling was evaluated using two AAK1 mutants: 1) AAK1-74A, which lacks kinase activity, and 2) AAK1-AID, which encodes a truncated form of AAK1 that cannot bind the AP2 complex^12^. Neither AAK1-74A nor AAK1-AID significantly repressed the BAR reporter (**Fig. 3b**), suggesting that both interaction and kinase activity are required for AAK1-mediated repression of WNT signaling. A role for clathrin and CME in AAK1-driven WNT suppression was tested by siRNA silencing of clathrin and by pharmacological inhibition clathrin-coated vesicles. Two non-overlapping clathrin siRNAs effectively silenced clathrin expression, and blocked AAK1-dependent suppression of the BAR reporter in HT1080 cells (**Fig. 3c**). Similarly, treatment with chlorpromazine, an established albeit promiscuous CME inhibitor^17,30,31^, suppressed WNT signaling induced by AAK1 silencing (Fig. 4D). These experiments suggest that AAK1 represses WNT signaling in a clathrin- and kinase-dependent manner.

**Figure 3.**
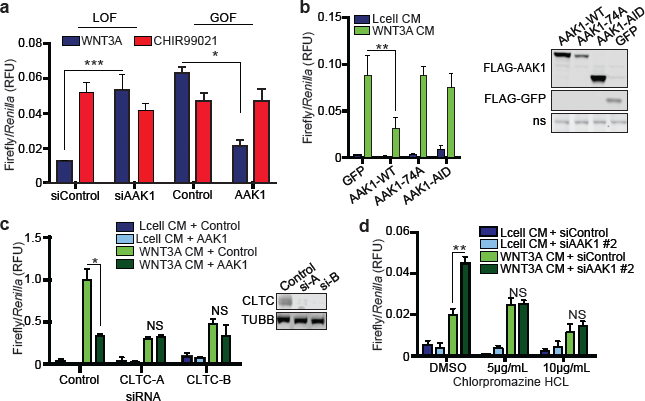
AAK1 represses β-catenin-mediated WNT signaling through regulation of endocytosis. (**a**) HT1080-B/R cells were transfected with indicated siRNA for 56 hours and then treated with WNT3A or CHIR99021 (1 µM) for 16 hours (left). HT1080-B/R cells were transfected with the indicated over-expression DNA construct and allowed to recover for 12 hours. Cells were then treated with WNT3A CM or CHIR99021 (1 ìM) for 16 hours (right). Data are averaged over 3 technical replicates. (**b**) BAR luciferase assay from HT1080 cells transiently transfected with BAR-firefly luciferase, TK-Ren, and the indicated expression constructs. Twelve hours post-transfection, cells were treated for 16 hours with Lcell or WNT3A CM and then assayed for BAR activity. Western blot of FLAG-AAK1 and FLAG-GFP is shown to illustrate expression of AAK1 constructs. Presented data is averaged across 3 technical replicates. (**c**) Luciferase assay in HT1080-B/R cells transfected with indicated siRNAs for 50 hours, then co-transfected with indicated expression constructs for 12 hours. Cells were then treated for 16 hours with either Lcell or WNT3A CM. Data are averaged over 3 technical replicates. Right panel shows Western blot of clathrin, indicating knockdown efficiency. (**d**) Luciferase assay of HT1080-B/R cells transfected with indicated siRNA for 56 hours, then treated as indicated for 16 hours. Data are averaged over 3 technical replicates. All panels: *** p<0.0005, ** p<0.005, * p<0.05. Data are representative of biological triplicates. For complete statistics, see methods and materials.

**Figure 4.**
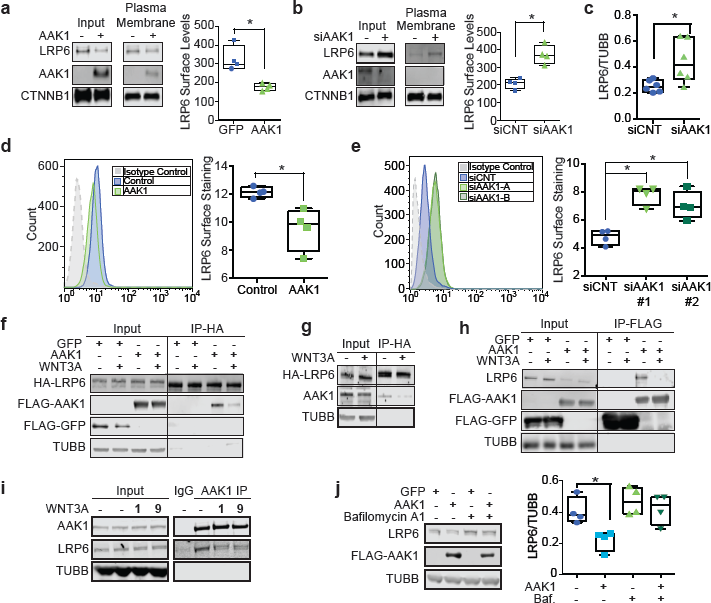
AAK1 represses LRP6 occupancy at the membrane. **(a,b)** Western blot analysis of HEK293T cells transfected with an AAK1 over-expression construct for 24 hours **(a)** or AAK1 siRNA for 72 hours **(b)** prior to membrane fractionation using a surface biotinylation assay. Data represent average surface staining of four biological replicate experiments quantified by LiCOR software. **(c)** LiCOR quantification of Western blot analysis of LRP6 levels with AAK1 knockdown. The box and whisker plot represents total LRP6 expression normalized to β-tubulin (loading control) from six biological replicates quantified by LiCOR software. **(d,e)** LRP6 surface staining as detected by flow cytometry analysis in HEK293T cells transfected with an AAK1 over-expression construct for 24 hours **(d)** or an AAK1 siRNA construct for 72 hours **(e)**. All experiments were quantified using FlowJo software. Experiment was performed in 4 biological replicates. **(f)** HEK293T cells were co-transfected with FLAG-AAK1 or control FLAG-hcRED and HA-LRP6 over-expression constructs. Twenty-four hours post-transfection, cells were treated with WNT3A CM for 1 hour. Western blot analysis of immunoprecipitated HA-LRP6. **(g)** HEK293T cells were transfected with HA-LRP6 for 24 hours and then treated with WNT3A CM for 1 hour. HA-LRP6 was immunoprecipitated, and the Western blot was probed for endogenous AAK1. **(h)** HEK293T cells were transfected with the FLAG-AAK1 or FLAG-GFP over-expression construct for 24 hours and then treated with WNT3A for 1 hour. IP of FLAG-AAK1 was conducted to visualize endogenous LRP6 as detected by Western blot. **(i)** Endogenous AAK1 IP in HEK293T cells treated as WNT3A CM for indicated times (untreated, 1, 9 hours) then endogenous LRP6 levels were visualized via Western blot analysis. **(j)** HEK293T cells were transfected with FLAG-AAK1 or FLAG-GFP over-expression constructs. Twenty-four hours post-transfection, cells were treated for 6 hours with the autophagy inhibitor, Bafilomycin A1 (100 nm) or DMSO control prior to cell harvest, then analyzed by Western blot for endogenous LRP6 and AAK1 levels. The box and whisker plot represents LRP6 expression normalized to β-tubulin from four biological replicates quantified by LiCOR software. All panels: Data are representative of biological triplicates, unless otherwise stated. * p<0.05. For complete statistical analysis, see methods and materials section.

We next examined the effect of AAK1 on LRP6 protein levels using cell surface biotinylation assays and flow cytometry. AAK1 over-expression decreased LRP6 protein levels at the plasma membrane (**Fig. 4a**). Additionally, AAK1 knockdown by siRNA increased LRP6 surface expression levels (**Fig. 4b**). Based on the observation that total LRP6 levels seem to decrease with AAK1 knockdown (**Fig. 4b**), we quantified total LRP6 expression across 6 biological replicates and demonstrate that with AAK1 knockdown, total LRP6 levels increase (**Fig. 4c**). To validate the changes in surface LRP6 expression findings in an orthogonal platform, we used flow cytometry to quantify cell surface LRP6 levels in HEK293T cells. Here, AAK1 overexpression decreased LRP6 surface levels while AAK1 knockdown increased LRP6 surface levels (**Fig. 4d,e**). Together, these data indicate that AAK1 negatively regulates LRP6 abundance on the plasma membrane. As such, we hypothesized that AAK1 and LRP6 co-complex. Using HEK293T cells, immunoprecipitation and Western blot analysis of HA-LRP6 protein complexes revealed the presence of over-expressed FLAG-AAK1 (**Fig. 4f**). Similarly, immunoprecipitation of HA-LRP6 co-purified endogenous AAK1 and reciprocally, immunopurification of over-expressed FLAG-AAK1 revealed co-complexed endogenous LRP6 (**Fig. 4g,h, respectively**). Finally, immunoprecipitation of endogenous AAK1 showed co-complexed endogenously expressed LRP6 (**Fig. 4i**). Interestingly, WNT3A stimulation suppressed the association of LRP6 and AAK1 in all experiments (**Fig. 4f-i**).

AAK1 silencing potentiated WNT3A-driven β-catenin activation and increased LRP6 protein levels in whole cell lysate and on the membrane (**Fig. 4b,c,e**). Because cargo targeted by clathrin-mediated endocytosis can be directed to lysosomes for degradation, we tested if the decreased LRP6 levels we observed following AAK1 over-expression was due to CME of LRP6 and subsequent lysosomal degradation. AAK1 was over-expressed in HEK293T cells before treatment with bafilomycin A1, an inhibitor of lysosomal acidification. Western blot analysis established that bafilomycin A1 treatment rescued AAK1-driven downregulation of LRP6 (**Fig. 4j**). Together, these results suggest that AAK1 inhibits WNT signaling by inducing CME endocytosis and lysosomal degradation of LRP6.

### WNT3A induces AAK1 phosphorylation of AP2M1

Within 15-30 minutes of WNT3A treatment, LRP6 is phosphorylated by GSK3β and CK1γ, resulting in transient suppression of β-catenin phosphorylation and degradation^3–6^.Extended duration of WNT3A stimulation of 4 to 6 hours results in multivesicular body formation and sequestration of GSK3β^23^. Here we asked if WNT3A treatment regulated AAK1-dependent phosphorylation of AP2M1. Time course analysis in HT1080 cells revealed WNT3A induced phosphorylation of AP2M1 at 8-10 hours post treatment, with no change in total AP2M1 or AAK1 protein levels (**Fig. 5a**). To rule out artifacts due to the use of WNT3A conditioned media and to extend relevance to a second cell type, we treated HT1080 cells (**Supplementary Fig. 3a**) and HEK293T (**Supplementary Fig. 3c**) cells with recombinant human WNT3A (rhWNT3A) over a time course of 4 to 16 hours. Western blot analysis revealed an increase in pAP2M1 protein levels 8-10 hours post WNT3A stimulation. Further, a pulse of WNT3A (15 minutes) induced a similar increase in AP2M1 phosphorylation as a prolonged WNT3A treatment (**Supplementary Fig. 3b, d**). The WNT3A-induced phosphorylation of AP2M1 required AAK1 expression and activity. First, HT1080 cells or HEK293T cells were treated with WNT3A CM and the AAK1 inhibitor, 25A, before Western blot analysis of phosphorylated AP2M1 — 25A blocked WNT3A-induced phosphorylation of AP2M1 (**Supplementary Fig. 3b,d**). Second, siRNA silencing of AAK1 similarly suppressed WNT3A-induced phosphorylation of AP2M1 (**Fig. 5b**). These data support a model where prolonged WNT3A treatment establishes an AAK1-driven negative feedback loop wherein CME of LRP6 decreases its abundance on the plasma membrane.

**Figure 5.**
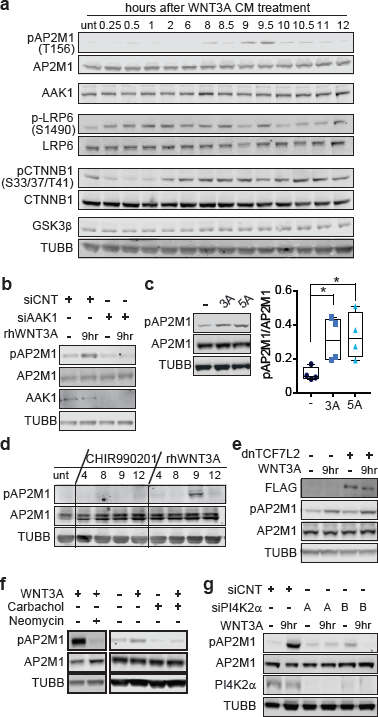
WNT treatment induces AAK1-, PIP2-dependent phosphorylation of AP2M1. **(a)** Western blot analysis of HT1080 cells treated with WNT3A CM for indicated times. **(b)** Western blot analysis of HT1080 cells transfected with either AAK1 or control siRNA for 72 hours. Cells were then treated with rhWNT3A (200 ng/mL) for 15 minutes, then the media was changed for fresh, complete DMEM and cells were incubated for 9 hours. **(c)** Western blot analysis of pAP2M1 levels in HEK293T cells treated with either rhWNT3A (3A) or rhWNT5A (5A) (200 ng/mL) for 9 hours. The box and whisker plot represents phosphorylated LRP6/total LRP6 expression from four biological replicates quantified by LiCOR software. **(d)** Western blot analysis of HT1080 cells treated with either CHIR990201 compound (1 µM) or rhWNT3A (200 ng/mL) for the indicated time. **(e)** HEK-293T cells were transfected with FLAG-dnTCF4 for 24 hours. Cells were then treated with WNT3A for 15 minutes, then the media was changed for fresh, complete DMEM, incubated for 9 hours, and analyzed by western blot. **(f)** Western blot analysis of HT1080 cells treated with WNT3A CM for 15 minutes, then the media was changed for complete DMEM containing either carbachol (100 µM) or neomycin (100 µM). Cells were incubated for 9 hours prior to cell harvest. **(g)** Western blot analysis of HT1080 cells transfected with siRNA against PI4K2α for 72 hours and pulsed with WNT3A CM for 15 minutes and then incubated for 9 hours. All experiments were performed in biological triplicate.

Intrigued with the delayed WNT3A-induced phosphorylation of AP2M1 by AAK1, we performed a series of functional assays to identify requisite signaling events. First, we found that AAK1-mediated pAP2M1 accumulation was induced by rhWNT3A (3A) and rhWNT5A (5A), the latter of which signals in a β-catenin-independent fashion (**Fig. 5c**). Second, inhibition of GSK3β with CHIR99021 did not result in elevated pAP2M1 levels, suggesting that WNT3A-induced pAP2M1 did not require β-catenin-dependent transcription (**Fig. 5d**). In agreement with these data, inhibiting WNT3A-dependent transcription with over-expressed dominant-negative TCF7L2/TCF4 (dnTCF7L2/dnTCF4) did not affect pAP2M1 accumulation (**Fig. 5e**). Third, previous studies established that PIP2 levels increase following short-term (4 hour) WNT3A treatment, and that PIP2 is required for LRP6 signalosome formation^19,21^. Further, PIP2 was shown to recruit AP2 to the signalosome, leading to clathrin-mediated endocytosis of LRP6^19^. Thus, we hypothesized that PIP2 is required for WNT3A-induced phosphorylation of AP2M1 by AAK1. To test this hypothesis, we inhibited PIP2 by treating cells with neomycin or carbachol, both of which block PIP2 signaling^32,33^. Treatment of HT1080 cells with neomycin or carbachol blocked WNT3A-induced phosphorylation of AP2M1 (**Fig. 5f**). Furthermore, we blocked PIP2 production by silencing the PI4K2α kinase with 2 non-overlapping siRNAs. PI4K2α phosphorylates phosphatidylinositol (PI) to phosphatidylinositol 4-phosphate (PI4P) and is required for subsequent PIP2 production^34^. In agreement with the neomycin or carbachol experiments, PI4K2α silencing in HT1080 cells blocked WNT3A-dependent increase in pAP2M1 (**Fig. 5g**). We conclude that WNT ligands drive AAK1-dependent phosphorylation of AP2M1 in a manner that is: 1) temporally-delayed from signalosome formation, 2) independent of β-catenin-mediated transcription and 3) dependent upon PIP2 production.

## Discussion

Clathrin-mediated endocytosis is a well-established regulator of WNT signaling, although whether CME promotes or inhibits the WNT pathway remains debated^7,17,19–21,23,24^. Certainly, contextual and temporal features in CME-directed WNT studies must be considered. As a central and positive regulator of CME, our discovery of AAK1 as a WNT modulator is of no surprise. Collectively our data show that AAK1 negatively regulates β-catenin-dependent WNT signaling through CME endocytosis of LRP6. Although the experiments have not yet been done, we speculate that Frizzled is similarly targeted by AAK1 and CME. In the course of our studies, we discovered an unexpected WNT-induced phosphorylation event on T156 of AP2M1. This was mediated by AAK1, and reproducibly occurred at 8-10 hours post WNT treatment. In Figure 8, we summarize our findings in a multi-step model that integrates WNT-driven signalosome formation, PIP2 production, AAK1, AP2M1 phosphorylation and CME of LRP6.

In the absence of WNT ligand, Frizzled and LRP6 remain dissociated, allowing for β-catenin phosphorylation, ubiquitylation and degradation (**Fig. 6a**)^3,4^. Within minutes of WNT stimulation, the LRP6 signalosome forms, resulting in transient repression of β-catenin phosphorylation by GSK3β (**Fig. 6b**)^7^. The newly formed LRP6 signalosome requires PIP2 for assembly and contains both clathrin and AP2 (**Fig. 6b**)^21^. Based on our data, we suggest that long-term (8-10 hours) WNT stimulation induces a spike in AAK1-dependent phosphorylation of T156-AP2M1. This promotes clathrin polymerization, clathrin-coated pit stabilization and ultimately CME of LRP6, thus constituting a WNT-driven and AAK1-dependent negative feedback loop.

**Figure 6.**
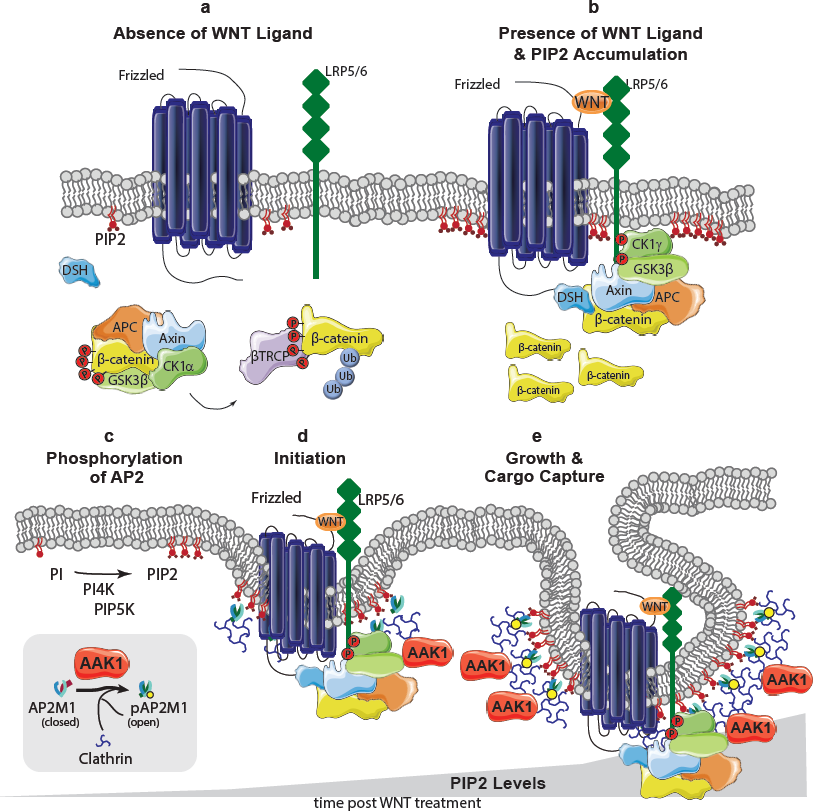
Proposed model for AAK1-dependent and clathrin-mediated endocytosis of LRP6. **(a)** In the absence of WNT ligand, Frizzled and LRP5/6 remain dissociated, allowing for the formation of the destruction complex, leading to the phosphorylation, ubiquitylation and degradation of β-catenin. PIP2 expression remains at low, basal levels. **(b)** In the presence of WNT ligand, Frizzled and LRP5/6 co-complex, allowing for recruitment of an alternate membrane associated complex, the LRP6 signalosome. This allows for the accumulation of β-catenin and activation of the WNT signaling pathway. Additionally, exposure to WNT ligand promotes the accumulation of PIP2. **(c)** Conversion of PI to PIP2 promotes an open conformation of AP2, potentiating clathrin binding. **(d)** We hypothesize that PIP2 levels continue to accumulate with longer WNT exposure. Phosphorylation of AP2M1 by AAK1 promotes binding to cargo sorting signals on LRP6, and enhances the growth and stabilization of clathrin-coated pits. **(e)** PIP2 levels accumulate to a specific threshold, driving AAK1 dependent AP2M1 phosphorylation. This results in growth of the clathrin-coated pit and cargo capture of the LRP6 signalosome, which undergoes CME and is ultimately degraded.

Phosphoinositide is converted to PIP2 by PI4K2α and PIP5K^34^. Importantly, PIP5K1 association with the DIX domain of DVL1/2/3 results in activation of PIP5K1^21,35^. We hypothesize that through this mechanism, WNT-driven recruitment of DVL to the LRP/Frizzled receptor complex results in a time-dependent increase in local PIP2 production. PIP2 and cargo proteins allosterically activate the AP2 complex, thus driving clathrin polymerization and maturation of clathrin-coated pits (**Fig. 6c**)^9,11^. Phosphorylation of AP2M1 by AAK1 stabilizes the AP2 complex in an open/active confirmation, promoting the binding of clathrin and cargo proteins and ultimately enhancing CME (**Fig. 6d**)^13^. Although we do not yet understand the mechanistic details, we speculate that PIP2 levels continue to rise hours after WNT stimulation, possibly triggering an increase in AAK1-dependent phosphorylation of AP2M1 and CME of LRP6 (**Fig. 6e**). Further studies that quantify PIP2 levels at 8-10 hours post WNT stimulation are needed. Alternatively, the AP2 complex has been reported to cycle between phosphorylated and dephosphorylated states during CME and that this cycling is necessary for robust CME^11^. Therefore, it is possible that WNT stimulation shifts the AAK1/phosphatase balance to favor phosphorylated AP2M1. The molecular events suggested in this time-delayed model are consistent with the timing reported in the GSK3-sequestration model proposed by De Robertis and colleagues^23^.

Central to our model is the increase in phosphorylated AP2M1 levels at 8-10 hours post-WNT treatment. We found that phosphorylation of T156 in AP2M1 was AAK1-, PIP2- and WNT ligand-dependent, but independent of β-catenin-driven transcription. Consistent with this, we also show that the β-catenin-independent WNT ligand, WNT5A, similarly induces pT156-AP2M1. The physiological significance of accumulated pT156-AP2M1 hours after WNT treatment remains unclear. Although our studies employed various cell models receiving bolus administration of WNT ligand, we hypothesize that in certain developmental or disease contexts, WNT-induced pT156-AP2M1 may establish a negative feedback loop to remove LRP/Frizzled from the cell membrane.

The AP2 complex binds Yxxɸ or dileucine motifs within the intracellular domains of transmembrane cargo proteins^36,37^. Through phosphorylation of T156-AP2M1, AAK1 was previously shown to promote association of the AP2 complex with cargo^10^. Interestingly, in coimmunoprecipitation experiments, we discovered an association between AAK1 and LRP6. AP2 interacts with LRP6 through a conserved Yxxɸ motif, which is required for LRP6 signalosome formation^19,24^. Thus, it is prossible that the AAK1-LRP6 co-complex reported here requires AP2 as a bridge. We also report that the AAK1-LRP6 co-complex dissolves following treatment with WNT3A. While further experimentation is needed to define the dynamics of protein-protein interactions during CME of LRP6, it is possible that following AAK1-dependent phosphorylation of AP2M1, AAK1 dissociates from the LRP6/AP2 complex.

AAK1 contributes to several neurological disorders, including neuropathic pain, Alzheimer’s Disease, Parkinson’s Disease, schizophrenia and Amyotrophic Lateral Sclerosis^38,39^.AAK1 has a described role in dendrite branching and spine development^40^, and through the regulation of Neuregulin1/ErbB4, AAK1 has been linked to schizophrenia^41^. Just recently, the LX9211 AAK1 inhibitor entered phase 1 clinical trials for neuropathic pain. Here we report the development of a potent and selective AAK1 pharmacological inhibitor (SGC-AAK1-1). SGCAAK1-1 demonstrates improved biochemical selectivity over the phase 1 clinical agent LX9211 and is confirmed to be cell active. Thus, we report the best available chemical tool to study AAK1/BMP2K pathways and related biology. Like the lead compound 25A, SGC-AAK1-1 inhibited AAK1-dependent phosphorylation of AP2M1 and activated WNT signaling. Looking forward, *in vitro* and *in vivo* neural-directed studies of SGC-AAK1-1 and its comparison with LX9211 are needed. Further, while we describe a role for AAK1 in negatively regulating WNT signaling, AAK1 also inhibits Neuregulin1/ErbB4 and positively regulates the NOTCH pathway^41,42^. Therefore, AAK1 inhibitors may also regulate these signaling cascades and consequently prove therapeutically beneficial in diseases with misregulated NOTCH or Neuregulin1/ErbB4 signaling.

## Materials and Methods

### Cell lines and tissue culture

All cells were cultured at 37 °C and 5% CO_2_. HT1080, HEK293T, RKO, Lcells, and WNT3A expressing Lcells were obtained from American Type Culture Collection (ATCC) (Manassas, VA) and grown in DMEM with 10% FBS. Each cell line was routinely tested for contamination and passaged no more than 20 passages from the original ATCC stock. Cells were treated at the indicated concentrations with the following compounds: CHIR99021 (Cayman Chemicals), chlorpromazine HCL (Sigma-Aldrich), bafilomycin A1 (Cayman Chemicals), rhWNT3A (PeproTech), rhWNT5A (R&D), neomycin trisulfate salt hydrate (Sigma-Aldrich) and carbachol (EMD Chemicals).

### WNT3A conditioned media

Conditioned media was collected as described by ATCC. Briefly, WNT3A and control Lcells were grown to 100% confluency before fresh media was added, conditioned for 48 hours, then collected.

### DNA constructs and siRNAs

The human kinase open reading frame library was obtained from Addgene (cat# 1000000014) and cloned into a custom pHAGE-CMV-FLAG vector using Gateway cloning technology. All constructs were N-terminally sequence verified. Wild type AAK1 and mutants were a gift from Dr. Sean Conner (University of Minnesota). GFP-LRP6, HALRP6, ANKRD6, and FLAG-dnTCF7L2 were kindly provided by Dr. Randell Moon (University of Washington). ADCK1, ADCK2, MAST1, and TGFBR3 were obtained as part of the kinome library from Addgene. All siRNAs were obtained from Life Technologies (ThermoFisher, Waltham, MA), and sequences are provided in Supplementary Table 5.

### Transcriptional reporter assays

All luciferase reporter assays and IncuCyte fluorescent reporter assays were performed as previously described^26^. Firefly luciferase and the *Renilla (Ren)* control were detected using the Promega Dual-Luciferase Reporter Assay System per the manufacturer’s protocol (Promega). Plates were read on the EnSpire plate reader from PerkinElmer. For IncuCyte fluorescent reporter assays, stable BAR-mCherry cells were treated and imaged as indicated using the IncuCyte Live Cell Analysis System from Essen BioScience. For loss-of-function assays, cell lines stably expressing the BAR-Firefly luciferase reporter and TK-*Ren* luciferase were used and transfected with RNAiMAX (Life technologies, ThermoFisher) for 72 hours. For gain of function studies, the BAR-reporter (20 ng), TK-*Ren* (10 ng), and indicated constructs (70 ng) were transfected with TansIT2020 (Mirus Bio) for 24 hours. Luciferase readouts were normalized using internal mCherry control and co-transfected TK-*Ren.*

### Real-Time Quantitative PCR

Quantitative PCR (qPCR) was performed as described previously^26^. Briefly, cells were treated as indicated and RNA was collected using PureLink RNA Mini Kit (Invitrogen). cDNA was generated from 1ug of RNA using the iScript cDNA Synthesis Kit (BioRad) following kit specifications. qPCR was performed using Fast SYBER Green Master Mix (Applied Biosystems) on the Applied Biosystems 7400HT following manufacturer specifications. Primers are listed in Supplementary Table 5, and were published previously^26^.

### Immunoprecipitation and immunoblotting

Immunoprecipitation experiments were performed as previously described^18^. Standard Western blotting techniques were utilized. All images were obtained using a LiCOR Odyssey imager and quantified with LiCOR software. For Western blot analysis, all antibodies were used at a concentration of 1:1000, with the exception of loading controls, which were used at 1:10,000. Antibodies used were as followed: Bethyl Labs (Montgomery, TX) – AAK1 (A302-146a); Cell Signaling (Danvers, MA) – CD44 (13113), Clathrin (4796), LRP6 (3395), phospho-LRP6 (S1490) (2568), phospho-AP2M1 (T156) (7399S),PIP4K2A (5527), phospho-CTNNB1 (S33/37/T41) (9561S); Sigma-Aldrich – FLAG M2 (F3165), HA (11867423001), and beta-tubulin (T8328); Abcam (Cambridge, UK) – LRP6 (ab75358), AP2M1 (ab7995), BD Biosciences (San Jose, CA) – CTNNB1 (163510), GSK3β (610201); Santa Cruz (Dallas, TX) – DVL3 (sc-8027). Secondary antibodies were used at 1:5000 dilution and purchased from LiCOR Biosciences. Specific antibodies used are as follows: IRDye^®^ 800CW Goat anti Mouse IgG (925-32210), IRDye^®^ 680LT Goat anti Mouse IgG (925-68020), IRDye^®^800CW Goat anti Rabbit IgG (925-32211), and IRDye^®^ 680LT Goat anti Rabbit (925-68021).

### Dose-response measurement of SGC-AAK1-1 inhibition of pAP2 formation

AP2M1 T156 phosphorylation levels were measured after 2 hour treatment with SGC-AAK1-1 at different concentrations or 10 µM 34A. HEK293T cells were plated in a 96-well plate (1.5 × 10^5^ cells/well) and the next day they were treated with the compounds for 2 hours, followed by protein extraction with RIPA buffer with protease and phosphatase inhibitors and analysis via SDS-PAGE. Antibodies used were as follows: AP2M1 (Invitrogen, #PA5-21360), p-AP2M1 (Thr156) (Abcam #ab109397) and β-actin (Santa Cruz #sc-47778). All antibodies were used at 1:1000 dilution.

### Surface staining of LRP6

HEK293T cell were transfected for 48 hours prior to splitting into 60 cm plates with 400,000 cells/plate and allowed to attach overnight. Cells were disassociated with 0.5 mM EDTA in DPBS and washed with FACS Buffer (2% FBS in DPBS). Cells were stained in FACS buffer for 1 hour at on ice using 5 μg/mL LRP6 antibody (R&D) or Isotype Control (R&D) and secondary staining with 1:100 anti-mouse PE (Jackson Immunoresearch) for 45 minutes. After staining, cells were fixed with 2% paraformaldehyde in FACS buffer and analyzed by the UNC-Flow Cytometry Core.

### Surface biotinylation

For **s**urface biotinylation assays, the Pierce Cell Surface Protein Isolation Kit (ThermoFisher) was utilized and manufacturer specifications were followed. Briefly, cells were grown to 70% confluency, washed 3 times with cold PBS, and then biotinylated for 30 minutes at 4 °C with NHS-SS-sulfo-linked biotin (0.25mg/mL). The free biotin was quenched, and then the samples were washed 3 times with cold TBS prior to lysis and sonication. Lysates were cleared and then incubated with Streptavidin beads (GE Healthcare) for 1 hour at 4 °C with nutation. Beads were washed 4 times with cold TBS and then proteins were eluted with LDS protein loading buffer supplemented with DTT at 95 °C for 10 minutes.

### Chemistry general procedures

Reagents were purchased from commercial suppliers and used without further purification. ^1^H and ^13^C NMR spectra were collected in methanol-*d*_4_ and recorded on Varian Inova 400 or Inova 500 spectrometers. Peak positions are given in parts per million (ppm) downfield from tetramethylsilane as the internal standard; J values are expressed in hertz. Thin-layer chromatography (TLC) was performed on silica gel F254 to evaluate reaction courses and mixtures. Flash chromatography was performed on Merck 60 silica gel (0.063–0.200 mm). All non-aqueous reactions were performed under nitrogen atmosphere. Solutions containing the final products were dried over Na_2_SO_4_ before filtration and concentration under reduced pressure using a rotary evaporator.

### N-(6-(3-((N,N-diethylsulfamoyl)amino)phenyl)-1H-indazol-3-yl)cyclopropanecarboxamide (SGC-AAK1-1)

3-Aminophenylboronic acid pinacol ester (1 eq., 50 mg, 0.228 mmol) in pyridine (1.2 mL) was cooled to 0 ^o^C in an ice bath. Diethylsulfamoyl chloride (2.06 eq., 80.7 mg, 0.0754 mL, 0.47 mmol) was added drop wise, and the reaction mixture was stirred overnight. The reaction progress was checked by TLC, revealing complete consumption of starting material (SM). Solvent was removed and the residue was partitioned between dichloromethane and saturated aqueous sodium bicarbonate solution. The organic phase was dried (Na_2_SO_4_), filtered and concentrated. The residue was purified using an Isco Combiflash companion automated purification system (4g (18 mL/min), SiO_2_, 80%/20% to 25%/75% heptanes/EtOAc) to give diethyl({[3-(tetramethyl-1,3,2-dioxaborolan-2-yl)phenyl]sulfamoyl})amine as an orange solid (44 mg, 54%), which was determined to be desired product by ^1^H NMR.

The following was added to a microwave vial: N-(6-bromo-1H-indazol-3-yl)cyclopropanecarboxamide (1 eq., 20 mg, 0.0677 mmol), diethyl({[3-(tetramethyl-1,3,2-dioxaborolan-2-yl)phenyl]sulfamoyl})amine (1 eq., 24 mg, 0.0677 mmol), and Pd(dppf)Cl2 (10%, 4.96 mg, 0.00677 mmol) in a mixture of dioxane (0.19 mL) and 1M aq sodium carbonate (0.16 mL). The vial was capped and heated in the microwave at 120 ^o^C for 30 minutes. Once cool, the reaction progress was examined by TLC, which revealed complete consumption of SM. The mixture was filtered through Celite and washed with EtOAc (ethyl acetate). The combined organic solution was washed with brine, dried (Na_2_SO_4_), filtered and concentrated. The residue was purified using an Isco Combiflash companion automated purification system (4g (18 mL/min), SiO_2_, 75%/25% to 20%/80% heptanes/EtOAc) to give **SGC-AAK1-1** as a yellow solid (14.3 mg, 49%): ^1^H NMR (400 MHz, Methanol-*d*_4_) δ 7.80 (d, *J* = 0.8 Hz, 1H), 7.58 (s, 1H), 7.48 (dt, *J* = 0.8, 2.1 Hz, 1H), 7.38 – 7.35 (m, 2H), 7.33 (d, *J* = 8.5 Hz, 1H), 7.17 – 7.12 (m, 1H), 3.29 (d, *J* = 1.6 Hz, 4H), 1.91 (tt, *J* = 4.5, 8.0 Hz, 1H), 1.05 (t, *J* = 7.1 Hz, 6H), 1.01 (dt, *J* = 3.0, 4.4 Hz, 2H), 0.95 – 0.88 (m, 2H); ^13^C NMR (101 MHz, Methanol-*d*_4_) δ 174.1, 142.1, 141.9, 139.8, 139.0, 129.1, 122.2, 121.6, 119.7, 118.3, 118.1, 107.5, 48.2, 41.6, 13.5, 12.5, 6.8.

### General procedure for preparation of 3-acylamino-6-bromoindazoles

To a solution of 6-bromo-1H-indazol-3-amine (48 mg, 0.23 mmol) in pyridine (0.75 mL) was added cyclopropanecarboxylic acid chloride (0.021 mL, 0.23 mmol, 1 eq.,) drop wise at 0 **°**C. The reaction mixture was stirred at this temperature for 4 hours and then allowed to warm to room temperature. Once the reaction was complete the solvent was removed under reduced pressure. The residue was then dissolved in N,N-dimethylformamide and water was added drop wise. The precipitated solid was then washed with hexanes 3 times and further dried to afford *N*-(6-bromo-1*H*-indazol-3-yl)cyclopropanecarboxamide (40.4 mg, 64% yield).

### General procedure for 3-acylamino-6-arylindazoles

The following was added to a microwave vial: *N*-(6-bromo-1H-indazol-3-yl)cyclopropanecarboxamide (344 mg, 1.12 mmol) dissolved in dioxane (4mL) and 1M Na_2_CO_3_ (1mL). To this solution, the following was added: 3-aminophenylboronic acid pinacol ester (269 mg, 1.12 mmol, 1.0 eq.,), Pd(dppf)Cl_2_ (100.3 mg, 0.123 mmol, 0.1eq,). The reaction was run in the microwave at 160 **°**C for 20 minutes, at which time SM was consumed. The reaction mixture was poured into water and extracted 3 times with EtOAc. The combined organic layers were dried over Na_2_SO_4_ and concentrated to dryness. The compound was then purified by flash chromatography to afford *N*-(6-(3-aminophenyl)-1*H-*indazol-3-yl)cyclopropanecarboxamide (100 mg, 28% yield).

### *N*-(6-(3-(cyclopropanesulfonamido)phenyl)-1*H*-indazol-3-yl)cyclopropanecarboxamide (25A)

To a solution of *N*-[6-(3-aminophenyl)-1H-indazol-3-yl]cyclopropanecarboxamide (67mg, 0.2292 mmol) in pyridine (1mL) cooled to 0 **°**C, cyclopropanesulfonyl chloride (1eq, 32.22mg, 0.2292 mmol) was added drop wise. The reaction mixture was stirred at 0 **°**C for 4 hours and warmed to room temperature. After verification by LC/MS that SM had been consumed, solvent was removed under reduced pressure. The compound was then purified by high pressure liquid chromatography (HPLC) from 10% to 100% ACN/water + 0.05% TFA to yield **25A** (44.5 mg, 49% yield). ^1^H NMR (400 MHz, Methanol-*d*_4_) δ 7.83 – 7.78 (d, *J* = 8.6 Hz, 1H), 7.61 – 7.54 (m, 2H), 7.46 – 7.30 (m, 3H), 7.29 – 7.23 (ddd, *J* = 7.7, 2.2, 1.3 Hz, 1H), 2.62 – 2.49 (tt, *J* = 8.0, 4.8 Hz, 1H), 1.95 – 1.84 (ddd, *J* = 12.5, 7.9, 4.2 Hz, 1H), 1.07 – 0.77 (m, 8H). ^13^C NMR (126 MHz, Methanol-*d*_4_) δ 5.8 (2C), 8.4 (2C), 15.0, 30.5, 30.8, 109.1, 117.2, 121.2, 121.3, 121.4, 123.1, 124.7, 130.8, 140.1, 141.2, 143.4 (2C), 143.8, 175.6. MS+ (– ES API) - 397.1.

### *N*-(6-(3-(cyclopropanesulfonamido)phenyl)-1*H*-indazol-3-yl)isobutyramide (SGC-AAK-1N)

^1^H NMR (400 MHz, Methanol-*d*_4_) δ 7.82 – 7.75 (d, *J* = 8.6 Hz, 1H), 7.62 – 7.54 (dt, *J* = 10.9, 1.5 Hz, 2H), 7.47 – 7.31 (m, 3H), 7.31 – 7.22 (ddd, *J* = 7.8, 2.2, 1.3 Hz, 1H), 2.84 – 2.69 (hept, *J* = 6.8 Hz, 1H), 2.61 – 2.50 (tt, *J* = 8.0, 4.8 Hz, 1H), 1.29 – 1.15 (d, *J* = 6.8 Hz, 6H), 1.14 – 0.82 (m, 4H). ^13^C NMR (126 MHz, Methanol-*d*_4_) δ 5.8 (2C), 20.0 (2C), 30.5, 36.3, 109.2, 117.4, 121.3, 121.4 (2C), 123.0, 124.7, 130.8, 140.1, 141.2, 141.3, 143.4, 143.8, 179.3. MS+ (– ES API) - 399.1.

### Cloning, protein expression and purification

For crystallization of BMP2K, a construct of BMP2K residues 38-345 (NCBI NP_942595) encompassing the kinase domain with surface mutations K320A and K321A in expression vector pNIC-Zb was used^43^. The construct was transformed into BL21(DE3) cells that co-express λ-phosphatase and three rare tRNAs (plasmid pACYC-LIC+). The cells were cultured in TB medium containing 50 µg/mL kanamycin and 35 µg/mL chloramphenicol at 37 °C with shaking until the OD_600_ reached ~3 and then cooled to 18°C for 1 hour. Isopropyl β-D-1-thiogalactopyranoside (IPTG) was added to a final concentration of 0.1 mM and the cultures were left overnight at 18 °C. The cells were collected by centrifugation then resuspended in 2x lysis buffer (100 mM HEPES buffer, pH 7.5, 1.0 M NaCl, 20 mM imidazole, 1.0 mM tris(2-carboxyethyl)phosphine (TCEP), Protease Inhibitor Cocktail Set VII (Calbiochem, 1/500 dilution) and flash-frozen in liquid nitrogen. Cells were lysed by sonication on ice. The resulting proteins were purified using Ni-Sepharose resin (GE Healthcare) and eluted stepwise in binding buffer with 300 mM imidazole. Removal of the hexahistidine tag was performed at 4 °C overnight using recombinant TEV protease. Protein was further purified using reverse affinity chromatography on Ni-Sepharose followed by gel filtration (Superdex 200 16/60, GE Healthcare). Protein in gel filtration buffer (25 mM HEPES, 500 mM NaCl, 0.5 mM TCEP, 5% [v/v] glycerol) was concentrated to 12 mg/mL (measured by UV absorbance, using the calculated molecular weight and estimated extinction coefficient, using a NanoDrop spectrophotometer (Thermo Scientific)) using 30 kDa molecular weight cut-off centrifugal concentrators (Millipore) at 4 °C.

### Protein crystallization

The AAK1 inhibitor (dissolved in 100 % DMSO) was added to the protein in a 3-fold molar excess and incubated on ice for approximately 30 minutes. The mixture was centrifuged at 14,000 rpm for 10 minutes at 4 °C prior to setting up 150 nL volume sitting drops at three ratios (2:1, 1:1, or 1:2 protein-inhibitor complex to reservoir solution). Crystallization experiments were performed at 20 °C. Crystals were cryoprotected in mother liquor supplemented with 20-25% glycerol before flash-freezing in liquid nitrogen for data collection. Diffraction data was collected at the Diamond Light Source. The best diffracting crystals grew in 10% (v/v) Broad MW PEG smear, 3.2 M MgCl_2_, 100 mM Hepes pH 7.0 ^44^. Data was collected at 100 K at the Diamond Synchrotron beamline I02. Data collection statistics can be found in Supplementary Table 4.

### Crystal structure determination

Diffraction data were integrated using XDS^45^ and scaled using AIMLESS from the CCP4 software suite^46^. Molecular replacement was performed with Phaser^47^ using BMP2K/AZD7762 (PDB: 4W9W)^43^ as a search model. Automated model building was performed with Buccaneer^48^. Automated refinement was performed in PHENIX^49^. Coot^50^ was used for manual model building and real space refinement. Structure validation was performed using MolProbity^51^. Structure factors and coordinates have been deposited in the PDB with code 5IKW.

### Binding-displacement assays

The TR-FRET ligand binding-displacement assays for AAK1, BMP2K, GAK and STK16 were performed as previously described^52^.

### Kinome screening

The KINOMEscan assay panel was measured at DiscoverX Corporation as previously described^53^. Data collection can be found in Supplementary Table 2.

### Isothermal Titration Calorimetry

AAK1 and BMP2K proteins were produced as previously described^43^. Isothermal titration calorimetry measurements were made on a Microcal VP-ITC instrument at 25 °C. For the interaction of AAK1 with SGC-AAK1-1, the compound was diluted to 22 µM in ITC buffer from a stock at 10 mM in DMSO and loaded directly into the cell. AAK1 was dialyzed at 4 °C overnight into ITC buffer (20 mM HEPES pH 7.5, 150 mM NaCl, 1 mM TCEP) and loaded into the ITC syringe at a final concentration of 218 µM. Following thermal equilibration, AAK1 was titrated into the cell using serial injections of 8 µL until saturation was observed in the thermogram. The same method was repeated for the BMP2K versus SGC-AAK1-1 interaction where protein was loaded into the syringe at a concentration of 288 µM and injected into a 32 µM solution of SGC-AAK1-1. The ITC data was analyzed with NITPIC^54^ and SEDPHAT^55^. The final fitted data values are in Supplementary Table 3.

### Measurement of *in vitro* IC50 values for inhibition of AAK1 enzymatic activity

Enzymatic activity of AAK1 was monitored in 20 µL reaction volumes containing 25 nM AAK1 (27-365) in 15 mM MOPS pH 7.5, 2 mM MgCl_2_ and 0.004% triton X-100 buffer plus 46 µM ATP (at *K*_m_) and 200 µM of an AP-2 derived peptide (biotin-GGSQITSQVTGQIGWRR-amide). Optimal activity was achieved with sample incubation at 37 ^o^C for 40 minutes (steady-state conditions) in buffer conditions that were tailored by factorial design using Design-Expert software (Stat-Ease, version 10). To generate inhibition curves, end-point reactions were setup having compounds pre-incubated in a reaction mix without ATP for 10 minutes at room temperature. Next, all samples (in PCR tubes) were transferred to a PCR instrument set at 37 ^o^C for another 5 minutes. The reaction started with the addition of 46 µM ATP (at *K*_m_), maintaining the incubation conditions for another 40 minutes. The reactions were stopped and the amount of ADP produced was measured using a coupled enzyme assay system that converts Amplex Red to Resorufin^56^. Fluorescence readings were made in a BMG Labtech Clariostar instrument with excitation peak set at 530 nm and emission peak reading at 590 nm.

### NanoBRET measurements

Constructs for NanoBRET measurements of AAK1 and BMP2K were kindly provided by Promega. The AAK1 construct represents the short isoform of AAK1 with an N-terminal Nanoluc fusion, and the BMP2K construct represents residues the short isoform of BMP2K with an N-terminal Nanoluc fusion. HEK293T cells (ATCC) were maintained in DMEM supplemented with 10% fetal bovine serum (FBS) (Life Technologies) with penicillin and streptomycin. NanoLuc-AAK1 or NanoLuc-BMP2K fusion constructs were complexed with Lipofectamine 2000 according to the manufacturer’s protocol (Invitrogen). DNA:Lipofectamine complexes were formed with 24 µg DNA and 60 µL Lipofectamine. The transfection complexes were then mixed with HEK293T cells in a 100 mm dish at 50-70% confluence in serum-free DMEM (Lonza), followed by incubation in a humidified, 37 °C, 5% CO_2_ incubator. After 4 hours of incubation, the medium was replaced by 10% FBS DMEM with antibiotics and the cells were incubated for an additional 20 hours. The NanoBRET assay was performed as previously described^57^. Briefly, cells were trypsinized and resuspended in Opti-MEM without phenol red (Life Technologies). Cells (85 µL) were then seeded into white, nonbinding surface plates (Corning) at a density of 2 × 10^4^ cells/well. Diluted tracer was prepared from 200 µM stock in Tracer Dilution Buffer (32.25% PEG400 in 12.5 mM HEPES Buffer pH 7.5) at a 1 to 4 ratio, and 5 µL was added to the cells to have a final concentration of 2, 1, 0.5 or 0.25 µM. All chemical inhibitors were prepared as concentrated stock solutions in dimethylsulfoxide (DMSO) (Sigma-Aldrich). Serial dilutions of the inhibitors at 50x the final assay concentration were made by dilution with DMSO, before the inhibitors were added to the cells at 1:50 ratio (2% final DMSO concentration). Cells were then incubated for 2 hours before Bioluminescence resonance energy transfer (BRET) measurements. To measure BRET, NanoBRET NanoGlo Substrate (Promega) was added, and filtered luminescence was measure on a BMG LABTECH Clariostar luminometer equipped with 450 nm BP filter (donor) and 610 nm LP filter (acceptor), using 0.5 s integration time with gain settings of 3,600 for both filters. Background-corrected BRET ratios were determined by subtracting the BRET ratios of samples with no tracer added.

### Statistics

All error bars are +/- standard error (SE). Statistical significance was evaluated by Mann-Whitney t-test unless otherwise stated. N values are stated in figure legends. Each panel was performed in at least biological triplicate, with higher replicates noted in figure legend. *** p<0.0005, ** p<0.005, * p<0.05.

## Acknowledgements

M.B.M. acknowledges support from the National Institutes of Health (RO1-CA187799 and U24-DK116204-01). M.J.A received financial support from National Institutes of Health T32 Predoctoral Training Grant in Pharmacology (T32-GM007040-43, T32-GM007040-42) and Initiative for Maximizing Student Diversity Grant (R25-GM055336-16). M.P.W. received support from the Lymphoma Research Foundation (337444) and the National Institutes of Health (T32-CA009156-35). The UNC Flow Cytometry Core Facility is supported in part by P30 CA016086 Cancer Center Core Support Grant to the UNC Lineberger Comprehensive Cancer Center. The SGC is a registered charity (number 1097737) that receives funds from AbbVie, Bayer Pharma AG, Boehringer Ingelheim, Canada Foundation for Innovation, Eshelman Institute for Innovation, Genome Canada, Innovative Medicines Initiative (EU/EFPIA) [ULTRA-DD grant no. 115766], Janssen, Merck & Co., Merck KGaA (Darmstadt, Germany), Novartis Pharma AG, Ontario Ministry of Economic Development and Innovation, Pfizer, São Paulo Research Foundation-FAPESP (2013/50724-5), Takeda, and Wellcome Trust [106169/ZZ14/Z]. R.R.R.S received financial support from FAPESP (2016/17469-0). The authors would like to thank members of the Major Laboratory for their feedback and expertise regarding experimental design and project direction. The authors also thank Claire Strain-Damerell and Pavel Savitsky for cloning various mutants of AAK1 and BMP2K proteins that were used in the crystallization trials.

## Author Contributions

M.J.A. co-led molecular characterization of AAK1, designed and carried out molecular biology experiments, and drafted and wrote manuscript. M.P.W. co-led molecular characterization of AAK1, designed and carried out molecular biology experiments and contributed to writing. A.D.A. synthesized SGC-AAK1-1, led SGC probe declaration effort, and contributed to writing. R.R.R. carried out NanoBRET measurements and cellular IC50 measurements (pAP2 Westerns). A.D.R. contributed to cloning and carrying out kinome gain-of-function screen and validation. D.M.G. contributed to AAK1 signaling characterization. M.G. carried out flow cytometry experiments. D.S.S. carried out dose response and pAP2M1 characterization of SGC-AAK1-1 and 25A. J.M.B. carried out NAK TR-FRET selectivity panel. R.M.C. made AAK1/BMP2K proteins and produced co-crystal structures. D.H.D. proposed and designed analogues. J.M.E. facilitated experimental design and contributed to writing. O.F. facilitated *in vitro* assays. C.G. made AAK1 proteins for crystallization/assays. O.G. facilitated protein synthesis and crystallization. P.H.G. carried out TR-FRET measurements and *in vitro* IC_50_measurements. N.K. executed compound scale-up for validation studies. S.M. contributed ideas and managed SGC probe declaration effort. A.S.S. made AAK1/BMP2K proteins for crystallization/assays. F.J.S. made proteins for NAK selectivity panel and ITC and carried out ITC experiments. C.I.W. synthesized 25A and SGC-AAK1-1N. T.M.W. managed chemistry efforts and aided in the design of molecules. W.J.Z. designed molecules, facilitated SGC probe declaration, and contributed to writing. M.B.M facilitated experimental design, project management and contributed to writing.

---

**Supplementary Figure 1.**
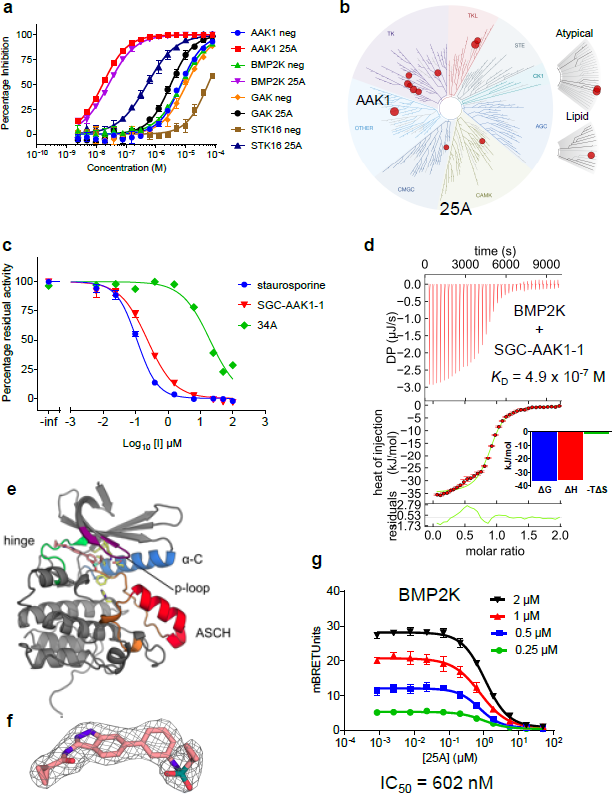
SGC-AAK1-1 is a potent and specific inhibitor of AAK1 and BMP2K that is active in cells. **(a)** Inhibitor 25A binds with high affinity to AAK1 and BMP2K but not to GAK and STK16 (related kinases from the same branch of the phylogenetic tree). Negative control compound 34A (illustrated as “neg”) does not bind to any of the kinases. The TR-FRET binding displacement assay was measured in duplicate. **(b)** Inhibitor 25A is most potent for AAK1 out of the 406 kinases tested and has good overall selectivity. **(c)** Inhibitor SGC-AAK1-1 inhibits AAK1 enzymatic activity in vitro with an IC_50_ of 233 nM at 46 µM ATP, similar to that of the potent but non-specific inhibitor staurosporine. Negative control compound 34A had an IC_50_ of 20 µM on AAK1. **(d)** SGC-AAK1-1 binds to BMP2K with weaker affinity than to AAK1, with a less favorable enthalpy of interaction. **(e)** The position of the inhibitor bound in the ATP binding site of BMP2K is shown. The Numb-associated-kinase-specific activation segment C-terminal helix (ASCH) is shown in red. **(f)** The inhibitor 25A is well resolved in the electron density. A 2Fo-Fc map is shown contoured at 1.0σ. **(g)** Inhibitor 25A binds to BMP2K in live HEK293T cells with an IC_50_ of 602 nM, derived from linear regression of the IC_50_ values at different concentrations of NanoBRET tracer. All experiments were performed in bioligical triplicates, unless otherwise noted. For complete statistical analysis, see methods and materials.

**Supplementary Figure 2.**
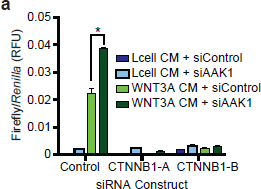
β-catenin knockdown impairs AAK1 regulation of WNT signaling. **(a)** Stable HT1080-B/R cells were transfected with the either β-catenin, AAK1 or control for 56 hours. Cells were then treated for 16 hours with WNT3A or Lcell CM. Bars represent average Firefly-/*Ren* (RFU) from 3 technical replicates +/-standard error (S.E.). Data are representative of biological triplicates. * p<0.05.

**Supplemental Figure 3.**
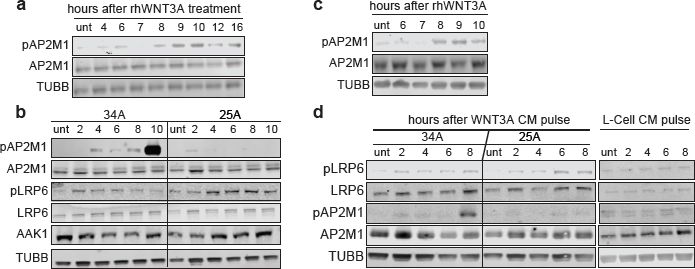
WNT treatment induces AAK1 mediated phosphorylation of AP2M1. **(a)** Western blot analysis of HT1080 cells treated with rhWNT3A (200 ng/mL) for indicated times. Data is representative of biological triplicates. **(b)** Western blot analysis of HT1080 cells stimulated with WNT3A CM for 15 minutes, then the media was replaced with complete DMEM. Cells were then treated with the AAK1 inhibitor, 25A, (1.25 µM) or small molecule analog, 34A, (1.25 µM) for the indicated times. **(c)** Western blot analysis of HEK293T cells treated with rhWNT3A (200 ng/mL) for indicated times. **(d)** Western blot analysis of HEK293T cells stimulated with WNT3A CM or Lcell CM for 15 minutes, then media was replaced for fresh, complete DMEM. Cells were then treated with the AAK1 inhibitor, 25A, (1.25 µM) or small molecule analog, 34A, (1.25 µM) for the indicated times. All experiments performed in biological triplicates.

**Supplementary Table 3.** Isothermal Titration Calorimetry data for binding of AAK1 or BMP2K proteins to SGC-AAK1-1.

|  | AAK1 | BMP2K |
| --- | --- | --- |
| $K_D$ (nM) | 117.5 | 487.5 |
| $\Delta H$ (kJ/mol) | -70.8 | -35.0 |
| $-T\Delta S$ (kJ/mol) | 31.2 | -1.0 |
| Fraction of incompetent ligand | 0.28 | 0.11 |

**Supplementary Table 4.** Data collection and refinement statistics for the BMP2K co-crystal structure with inhibitor 25A.

|  |  |
| --- | --- |
| PDB ID | 5IKW |
| Space group | *I*222 |
| Unit cell dimensions<br>*a*, *b*, *c* (Å), $\alpha$, $\beta$, $\gamma$ (°) | 42.0, 111.2, 164.1,<br>90, 90, 90 |
| **Data collection** |  |
| Resolution range (Å)<sup>a</sup> | 28.6-2.4<br>(2.5-2.4) |
| Unique observations<sup>a</sup> | 81078 (7855) |
| Average multiplicity<sup>a</sup> | 5.3 (5.2) |
| Completeness (%)<sup>a</sup> | 99.1 (95.2) |
| $R_{\text{merge}}$<sup>a</sup> | 0.074 (0.49) |
| Mean ($\langle I \rangle / \sigma(I)$)<sup>a</sup> | 13.2 (3.1) |
| Mean CC(1/2) | 0.991 (0.482) |
| **Refinement** |  |
| Resolution range (Å) | 21.75-2.41 |
| $R$-value, $R_{\text{free}}$ | 0.18, 0.22 |
| r.m.s. deviation from ideal<br>bond length (Å) | 0.010 |
| r.m.s. deviation from ideal<br>bond angle (°) | 1.04 |
| Ramachandran Outliers | 0 |
| Favoured | 98.3% |

<sup>a</sup> Values within parentheses refer to the highest resolution shell.

**Supplementary Table 5.**
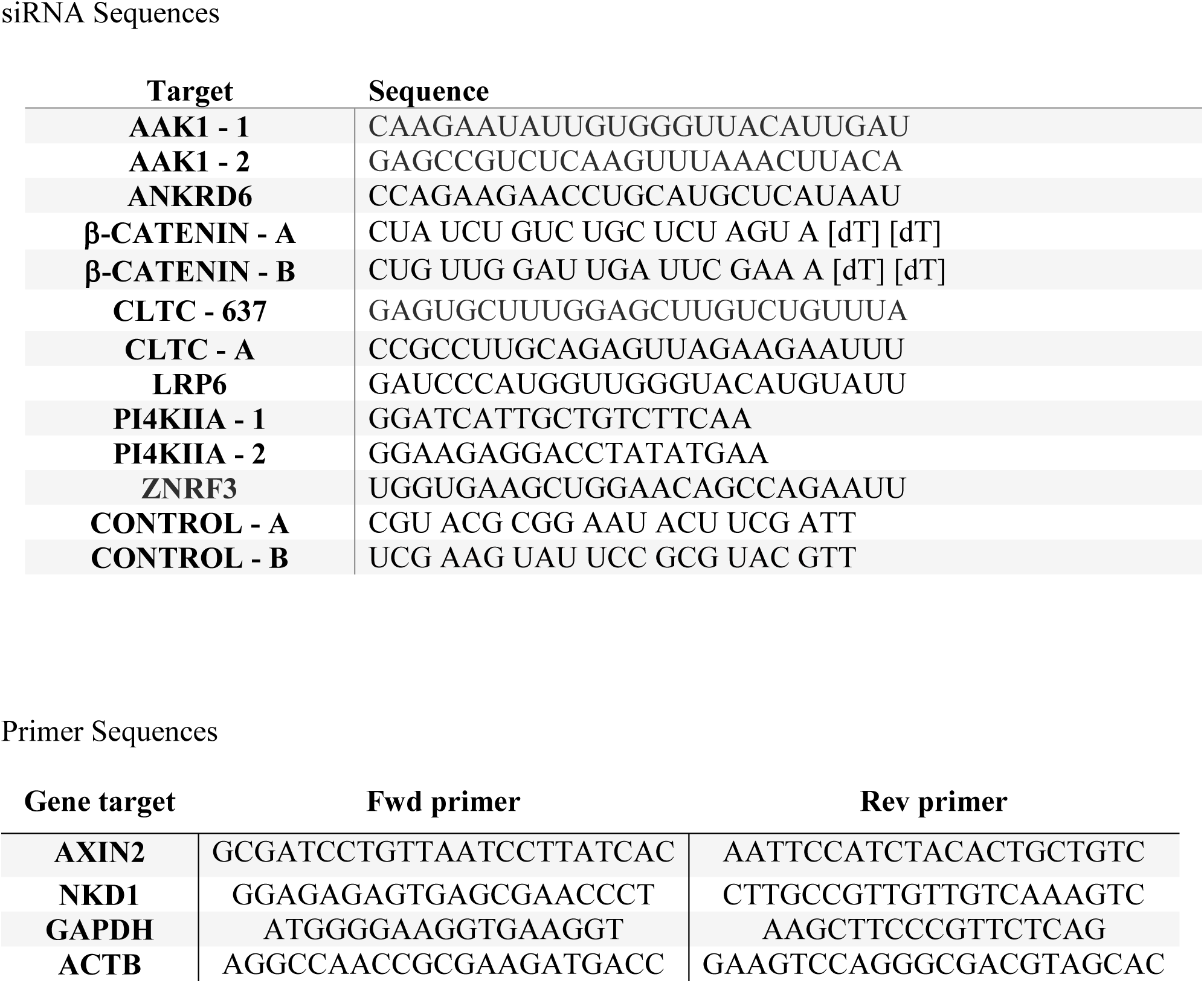
siRNA and primer sequences.

## References

1 Nusse R. & Clevers H. Wnt/beta-Catenin Signaling, Disease, and Emerging Therapeutic Modalities. Cell 169, 985–999, doi:10.1016/j.cell.2017.05.016 (2017).

2 Angers S. & Moon R. T. Proximal events in Wnt signal transduction. Nat Rev Mol Cell Biol 10, 468–477, doi:10.1038/nrm2717 (2009).

3 Liu C. et al. Control of beta-catenin phosphorylation/degradation by a dual-kinase mechanism. Cell 108, 837–847 (2002).

4 Liu C. et al. beta-Trcp couples beta-catenin phosphorylation-degradation and regulates Xenopus axis formation. Proc Natl Acad Sci U S A 96, 6273–6278 (1999).

5 Zeng X. et al. A dual-kinase mechanism for Wnt co-receptor phosphorylation and activation. Nature 438, 873–877, doi:10.1038/nature04185 (2005).

6 Hernandez A. R., Klein A. M. & Kirschner M. W. Kinetic responses of beta-catenin specify the sites of Wnt control. Science 338, 1337–1340, doi:10.1126/science.1228734 (2012).

7 Bilic J. et al. Wnt induces LRP6 signalosomes and promotes dishevelled-dependent LRP6 phosphorylation. Science 316, 1619–1622, doi:10.1126/science.1137065 (2007).

8 Behrens J. et al. Functional interaction of beta-catenin with the transcription factor LEF-1. Nature 382, 638–642, doi:10.1038/382638a0 (1996).

9 Honing S. et al. Phosphatidylinositol-(4,5)-bisphosphate regulates sorting signal recognition by the clathrin-associated adaptor complex AP2. Mol Cell 18, 519–531, doi:10.1016/j.molcel.2005.04.019 (2005).

10 Ricotta D., Conner S. D., Schmid S. L., von Figura K. & Honing S. Phosphorylation of the AP2 mu subunit by AAK1 mediates high affinity binding to membrane protein sorting signals. J Cell Biol 156, 791–795, doi:10.1083/jcb.200111068 (2002).

11 Kadlecova Z. et al. Regulation of clathrin-mediated endocytosis by hierarchical allosteric activation of AP2. J Cell Biol 216, 167–179, doi:10.1083/jcb.201608071 (2017).

12 Conner S. D. & Schmid S. L. Identification of an adaptor-associated kinase, AAK1, as a regulator of clathrin-mediated endocytosis. J Cell Biol 156, 921–929, doi:10.1083/jcb.200108123 (2002).

13 Conner S. D., Schroter T. & Schmid S. L. AAK1-mediated micro2 phosphorylation is stimulated by assembled clathrin. Traffic 4, 885–890 (2003).

14 Henderson D. M. & Conner S. D. A novel AAK1 splice variant functions at multiple steps of the endocytic pathway. Mol Biol Cell 18, 2698–2706, doi:10.1091/mbc.E06-09-0831 (2007).

15 Damke H., Baba T., Warnock D. E. & Schmid S. L. Induction of mutant dynamin specifically blocks endocytic coated vesicle formation. J Cell Biol 127, 915–934 (1994).

16 Hagemann A. I. et al. In vivo analysis of formation and endocytosis of the Wnt/beta-catenin signaling complex in zebrafish embryos. J Cell Sci 127, 3970–3982, doi:10.1242/jcs.148767 (2014).

17 Blitzer J. T. & Nusse R. A critical role for endocytosis in Wnt signaling. BMC Cell Biol 7, 28, doi:10.1186/1471-2121-7-28 (2006).

18 Jiang Y., He X. & Howe P. H. Disabled-2 (Dab2) inhibits Wnt/beta-catenin signalling by binding LRP6 and promoting its internalization through clathrin. EMBO J 31, 2336–2349, doi:10.1038/emboj.2012.83 (2012).

19 Kim I. et al. Clathrin and AP2 are required for PtdIns(4,5)P2-mediated formation of LRP6 signalosomes. J Cell Biol 200, 419–428, doi:10.1083/jcb.201206096 (2013).

20 Yamamoto H., Sakane H., Yamamoto H., Michiue T. & Kikuchi A. Wnt3a and Dkk1 regulate distinct internalization pathways of LRP6 to tune the activation of beta-catenin signaling. Dev Cell 15, 37–48, doi:10.1016/j.devcel.2008.04.015 (2008).

21 Pan W. et al. Wnt3a-mediated formation of phosphatidylinositol 4,5-bisphosphate regulates LRP6 phosphorylation. Science 321, 1350–1353, doi:10.1126/science.1160741 (2008).

22 Liu C. C., Kanekiyo T., Roth B. & Bu G. Tyrosine-based signal mediates LRP6 receptor endocytosis and desensitization of Wnt/beta-catenin pathway signaling. J Biol Chem 289, 27562–27570, doi:10.1074/jbc.M113.533927 (2014).

23 Taelman V. F. et al. Wnt signaling requires sequestration of glycogen synthase kinase 3 inside multivesicular endosomes. Cell 143, 1136–1148, doi:10.1016/j.cell.2010.11.034 (2010).

24 Gammons M. V., Renko M., Johnson C. M., Rutherford T. J. & Bienz M. Wnt Signalosome Assembly by DEP Domain Swapping of Dishevelled. Mol Cell 64, 92–104, doi:10.1016/j.molcel.2016.08.026 (2016).

25 Madan B. et al. USP6 oncogene promotes Wnt signaling by deubiquitylating Frizzleds. Proc Natl Acad Sci U S A 113, E2945–2954, doi:10.1073/pnas.1605691113 (2016).

26 Walker M. P. et al. FOXP1 potentiates Wnt/beta-catenin signaling in diffuse large B cell lymphoma. Sci Signal 8, ra12, doi:10.1126/scisignal.2005654 (2015).

27 Biechele T. L. et al. Wnt/beta-catenin signaling and AXIN1 regulate apoptosis triggered by inhibition of the mutant kinase BRAFV600E in human melanoma. Sci Signal 5, ra3, doi:10.1126/scisignal.2002274 (2012).

28 Elkins J. M. et al. Comprehensive characterization of the Published Kinase Inhibitor Set. Nat Biotechnol 34, 95–103, doi:10.1038/nbt.3374 (2016).

29 Vasta J. D. et al. Quantitative, Wide-Spectrum Kinase Profiling in Live Cells for Assessing the Effect of Cellular ATP on Target Engagement. Cell Chem Biol, doi:10.1016/j.chembiol.2017.10.010 (2017).

30 Wang L. H., Rothberg K. G. & Anderson R. G. Mis-assembly of clathrin lattices on endosomes reveals a regulatory switch for coated pit formation. J Cell Biol 123, 1107–1117 (1993).

31 Phonphok Y. & Rosenthal K. S. Stabilization of clathrin coated vesicles by amantadine, tromantadine and other hydrophobic amines. FEBS Lett 281, 188–190 (1991).

32 Huang Y., Qureshi I. A. & Chen H. Effects of phosphatidylinositol 4,5-bisphosphate and neomycin on phospholipase D: kinetic studies. Mol Cell Biochem 197, 195–201 (1999).

33 Baron C. B., Pompeo J., Blackman D. & Coburn R. F. Common phosphatidylinositol 4,5-bisphosphate pools are involved in carbachol and serotonin activation of tracheal smooth muscle. J Pharmacol Exp Ther 266, 8–15 (1993).

34 Toker A. The synthesis and cellular roles of phosphatidylinositol 4,5-bisphosphate. Curr Opin Cell Biol 10, 254–261 (1998).

35 Hu J. et al. Resolution of structure of PIP5K1A reveals molecular mechanism for its regulation by dimerization and dishevelled. Nat Commun 6, 8205, doi:10.1038/ncomms9205 (2015).

36 Chen W. J., Goldstein J. L. & Brown M. S. NPXY, a sequence often found in cytoplasmic tails, is required for coated pit-mediated internalization of the low density lipoprotein receptor. J Biol Chem 265, 3116–3123 (1990).

37 Collawn J. F. et al. Transferrin receptor internalization sequence YXRF implicates a tight turn as the structural recognition motif for endocytosis. Cell 63, 1061–1072 (1990).

38 Kostich W. et al. Inhibition of AAK1 Kinase as a Novel Therapeutic Approach to Treat Neuropathic Pain. J Pharmacol Exp Ther 358, 371–386, doi:10.1124/jpet.116.235333 (2016).

39 Shi B., Conner S. D. & Liu J. Dysfunction of endocytic kinase AAK1 in ALS. Int J Mol Sci 15, 22918–22932, doi:10.3390/ijms151222918 (2014).

40 Ultanir S. K. et al. Chemical genetic identification of NDR1/2 kinase substrates AAK1 and Rabin8 Uncovers their roles in dendrite arborization and spine development. Neuron 73, 1127–1142, doi:10.1016/j.neuron.2012.01.019 (2012).

41 Kuai L. et al. AAK1 identified as an inhibitor of neuregulin-1/ErbB4-dependent neurotrophic factor signaling using integrative chemical genomics and proteomics. Chem Biol 18, 891–906, doi:10.1016/j.chembiol.2011.03.017 (2011).

42 Gupta-Rossi N. et al. The adaptor-associated kinase 1, AAK1, is a positive regulator of the Notch pathway. J Biol Chem 286, 18720–18730, doi:10.1074/jbc.M110.190769 (2011).

43 Sorrell F. J., Szklarz M., Abdul Azeez K. R., Elkins J. M. & Knapp S. Family-wide Structural Analysis of Human Numb-Associated Protein Kinases. Structure 24, 401–411, doi:10.1016/j.str.2015.12.015 (2016).

44 Chaikuad A., Knapp S. & von Delft F. Defined PEG smears as an alternative approach to enhance the search for crystallization conditions and crystal-quality improvement in reduced screens. Acta Crystallogr D Biol Crystallogr 71, 1627–1639, doi:10.1107/S1399004715007968 (2015).

45 Kabsch W. Xds. Acta Crystallogr D Biol Crystallogr 66, 125–132, doi:10.1107/S0907444909047337 (2010).

46 Winn M. D. et al. Overview of the CCP4 suite and current developments. Acta Crystallogr D Biol Crystallogr 67, 235–242, doi:10.1107/S0907444910045749 (2011).

47 McCoy A. J. et al. Phaser crystallographic software. J Appl Crystallogr 40, 658–674, doi:10.1107/S0021889807021206 (2007).

48 Cowtan K. The Buccaneer software for automated model building. 1. Tracing protein chains. Acta Crystallogr D Biol Crystallogr 62, 1002–1011, doi:10.1107/S0907444906022116 (2006).

49 Adams P. D. et al. PHENIX: a comprehensive Python-based system for macromolecular structure solution. Acta Crystallogr D Biol Crystallogr 66, 213–221, doi:10.1107/S0907444909052925 (2010).

50 Emsley P., Lohkamp B., Scott W. G. & Cowtan K. Features and development of Coot. Acta Crystallogr D Biol Crystallogr 66, 486–501, doi:10.1107/S0907444910007493 (2010).

51 Chen V. B. et al. MolProbity: all-atom structure validation for macromolecular crystallography. Acta Crystallogr D Biol Crystallogr 66, 12–21, doi:10.1107/S0907444909042073 (2010).

52 Asquith C. R. M. et al. Identification and Optimization of 4-Anilinoquinolines as Inhibitors of Cyclin G Associated Kinase. ChemMedChem 13, 48–66, doi:10.1002/cmdc.201700663 (2018).

53 Davis M. I. et al. Comprehensive analysis of kinase inhibitor selectivity. Nat Biotechnol 29, 1046–1051, doi:10.1038/nbt.1990 (2011).

54 Keller S. et al. High-precision isothermal titration calorimetry with automated peak-shape analysis. Anal Chem 84, 5066–5073, doi:10.1021/ac3007522 (2012).

55 Zhao H., Piszczek G. & Schuck P. SEDPHAT—a platform for global ITC analysis and global multi-method analysis of molecular interactions. Methods 76, 137–148, doi:10.1016/j.ymeth.2014.11.012 (2015).

56 Charter N. W., Kauffman L., Singh R. & Eglen R. M. A generic, homogenous method for measuring kinase and inhibitor activity via adenosine 5′-diphosphate accumulation. J Biomol Screen 11, 390–399, doi:10.1177/1087057106286829 (2006).

57 Robers M. B. et al. Target engagement and drug residence time can be observed in living cells with BRET. Nat Commun 6, 10091, doi:10.1038/ncomms10091 (2015).

